# CERIUM OXIDE NANOPARTICLES ATTENUATE THE PRO-ARRHYTHMIC EFFECT OF DIESEL EXHAUST PARTICLES IN ISOLATED RAT HEARTS BY REDUCING OXIDATIVE STRESS

**DOI:** 10.1101/2025.04.11.648481

**Authors:** Freddy G. Ganse, Lena M. Ernst, Cristina Rodríguez, Marisol Ruiz-Meana, Javier Inserte, José Martínez-González, Ana Briones, Ana Belén García-Redondo, Marta Consegal, Elisabet Miró-Casas, Laia Yáñez-Bisbe, Aitor Pomposo, Marta Prades-Martínez, Ignacio Ferreira-González, Victor Puntes, Begoña Benito, Antonio Rodríguez-Sinovas

**Author notes:** **Co-senior and Co-corresponding authors:** Dr. Antonio Rodríguez-Sinovas and Dr. Begoña Benito, Cardiovascular Diseases Research Group, Vall d’Hebron University Hospital and Research Institute, Universitat Autònoma de Barcelona (Departament de Medicina), Pg. Vall d’Hebron 119, 08035 Barcelona, Spain.; Dr. Víctor Puntes, Disseny i Farmacodinàmica de Nanopartícules, Vall d’Hebron Research Institute, Barcelona, Spain; Catalan Institute of Nanoscience and Nanotechnology (ICN2), CSIC and BIST, Campus UAB, Bellaterra, 08193 Barcelona, Spain; Institució Catalana de Recerca I Estudis Avançats (ICREA), 08010 Barcelona, Spain.

## Abstract

**Introduction:** Epidemiological studies suggest an association between air pollution and ventricular arrhythmias, with reactive oxygen species (ROS) playing a crucial role. However, the causal relationship and long-term effects remain uncertain, and the effectiveness of interventions aimed at reducing ROS requires further investigation.

**Aims:** To evaluate the effects of chronic exposure to diesel exhaust particles (DEPs) on ventricular arrhythmogenesis, explore the underlying mechanisms, and assess the potential of cerium oxide nanoparticles (CeO_2_NP) as a ROS-detoxifying intervention.

**Methods:** Sprague-Dawley rats underwent intratracheal instillation of saline without or with DEPs (7.5 g/Kg for 1-3 weeks). Ventricular arrhythmia inducibility was then assessed in isolated hearts using a protocol of programmed electrical stimulation. Cardiac hypertrophy, collagen content, inflammation and oxidative stress were analyzed using histology, Western blot, RT-PCR, and measurement of malondialdehyde content. The potential protective effects of CeO_2_NP (0.5 mg/Kg/week, i.p.) were also tested.

**Results:** DEP exposure for 3 weeks increased the incidence and duration of sustained ventricular tachyarrhythmias (VTs), a finding that correlated with a moderate increase in interstitial collagen (from 3.11±0.12% in controls to 4.80±0.21% in DEP-exposed rats, p<0.001), and an early upregulation in the expression of collagen and other fibrotic and inflammatory markers. These effects associated with prolonged QRS complex and QTc intervals, and enhanced malondialdehyde content (356.7±21.2 vs. 455.3±17.2 μmol/g tissue, p=0.0066) after 3 weeks. CeO_2_NP treatment reduced oxidative stress and myocardial fibrosis, reversed electrocardiographic changes and attenuated DEP-induced pro-arrhythmic effects.

**Conclusion:** DEP exposure increases the incidence and duration of sustained VTs, collagen deposition and oxidative stress in rats. Treatment with CeO_2_NP attenuate these effects, arising as a potential novel strategy to mitigate the deleterious effects of air pollution.

## INTRODUCTION

The triad of pollution, climate change, and biodiversity loss constitutes the most pressing global environmental challenge of our time. Among these, pollution raises special concern due to its profound and widespread effects on human health. While recent years have seen reductions in deaths caused by household air pollution and water contamination, these gains have been overshadowed by rising fatalities linked to ambient air pollution and toxic chemical contamination ^1^. Deaths attributed to these modern pollution risk factors, unintended consequences of industrialization, have risen by 7% since 2015 and by over 66% since 2000 ^1^. Epidemiological evidence further identifies air pollution as one of the leading risk factors for mortality and disability for both males and females worldwide, with significant variations across different geographical regions ^2,3^.

Air pollution is composed of a complex mixture of gases (such as ground-level ozone, carbon monoxide, sulfur dioxide, and nitrogen oxides), semi-volatile liquids (methane, benzene, and polyaromatic hydrocarbons) and solid particles (aerosolized soil and dusts and other suspended particles, collectively referred to as particulate matter (PM)) that can be emitted by both natural (forest fires, volcanic eruptions, etc.) and anthropogenic (e.g., industry, traffic, household cooking, etc.) sources ^4^. Among these components, PM stands out, given its high toxicity and widespread distribution. PMs are formed by an elemental carbon core surrounded by a variety of chemicals, including sulphates and nitrates, redox-active metals, adsorbed soluble and vaporous hydrocarbons and organic carbon species. Based on their size, PM are categorized into coarse (PM10, 2.5-10 μm), fine (PM2.5, <2.5 μm), and ultrafine (PM0.1, <0.1 μm) particles ^4^.

In Europe, over 790,000 deaths annually can be attributed to air pollution, with about 48% of them being of cardiovascular origin ^5^. Indeed, clinical and epidemiological evidence indicates that both short- ^6–8^ and long-term ^8–10^ exposure to air pollution, and particularly to PM, is associated with increased cardiovascular mortality. Moreover, PM levels have been linked to a higher incidence of specific cardiovascular disorders, including heart failure ^11,12^, myocardial infarction ^6,11^, hypertension ^13^, aortic dissection ^14^ or stroke ^15^. In contrast to these relatively well-characterized adverse cardiovascular effects of air pollution, the relationship between air pollution and cardiac arrhythmias remains uncertain ^8^, with studies yielding conflicting results. Some works have failed to detect significant associations between exposure to air pollution and atrial or ventricular tachyarrhythmias ^16,17^, whereas others have reported that both short- ^18–20^ and long-term ^21–23^ exposure to PM increases the likelihood of cardiac arrhythmic events.

Experimental studies investigating the effects of air pollution on cardiac arrhythmias in animal models are relatively scarce. Most studies have focused on short-term exposure to diesel exhaust particles (DEPs), rich in PM, in animals with heightened susceptibility. These include models involving coronary artery occlusion ^24^, isoproterenol-induced heart failure ^25^ or in hypertensive rats in which arrhythmic events were triggered with aconitate ^26,27^. Similarly, airborne PM have been shown to induce arrhythmia-like cardiotoxicity in zebrafish embryos by altering the expression of several cardiac ion channels ^28^. These effects are associated with prolonged QT intervals and transmembrane action potentials, as well as the activation of CaMKII and the production of reactive oxygen species (ROS) ^29,30^.

Given the inconsistencies in clinical and epidemiological studies linking air pollution to cardiac arrhythmias and the limited scope of experimental research, lacking a clear cause-effect relationship and preventive strategies demonstrated thus far, we developed an experimental model of DEP exposure in rats with the following aims: (1) to analyze the effects of chronic intratracheal instillation of DEPs on ventricular arrhythmogenesis; and (2) to investigate the underlying mechanisms. Additionally, and as previous studies have linked the effects of air pollution to systemic and local inflammation as well as oxidative stress ^8,31,32^, we also investigated whether the effects of DEP exposure observed in our study were associated with this phenomena. In this context, rare-earth cerium oxide nanoparticles (CeO_2_NP) have emerged as safe and potent antioxidant and antiinflammatory agents ^33^. Cerium oxide nanoparticles, or nanoceria, exhibit defects in their crystal structure, resulting in the coexistence of both Ce^3+^ and Ce^4+^ ions ^33^. This unique property grants CeO_2_NP excellent catalytic activity and ROS-buffering capacity, mimicking the function of enzymes such as superoxide dismutase, catalase and peroxidase ^33^. Consequently, our third objective was to assess the impact of CeO_2_NP as a ROS-detoxifying strategy.

## METHODS

The data supporting the conclusions of this study are available from the corresponding authors. A complete description of the methods used in this work can be found in the Supplemental material. Animal studies complied with European legislation (Directive 2010/63/EU) on the protection of animals used for scientific purposes, with the Guide for the Care and Use of Laboratory Animals published by the US National Institutes of Health (NIH Publication No. 85-23, revised 1996, updated in 2011), and were approved by the Ethics Committee of Vall d’Hebron Research Institute (protocol number CEEA49.20, CEA/11228/P1/1).

### Preparation of diesel exhaust particles (DEPs)

DEPs (SRM-2975; National Institute of Standards and Technology, Gaithersburg, USA) were suspended in 0.9% sterile saline at a stock concentration of 20 mg/mL, vortexed and sonicated to minimize particle aggregation.

### Preparation of CeO_2_NP

#### Synthesis of CeO_2_NP

Cerium (III) chloride heptahydrate (10 mM) was precipitated with tetramethylammonium hydroxide (27 mM) in the presence of sodium citrate (20 mM). The mixture was stirred overnight at room temperature and then refluxed at 100°C for 4 hours, yielding 3 nm CeO_2_ NPs (1.72 mg/mL). CeO_2_NPs were purified using 3 kDa centrifugal filters (Amicon-Ultra-15 3K, Merck) and resuspended in 2.2 mM sodium citrate.

#### CeO_2_NP-RSA Conjugation

To enhance biocompatibility, CeO_2_NPs were conjugated with rat serum albumin (RSA, Merck) in 10 mM phosphate buffer (pH 7.4) at 4°C for 24 hours prior to injection.

#### Bacterial Endotoxin (LAL) test

All synthesis and purification steps were performed under sterile conditions with non-pyrogenic materials. Quantitative determination of lipopolysaccharide levels was conducted by Echevarne Analysis Laboratory (Barcelona, Spain).

#### Characterization of CeO_2_NPs

##### High-Resolution Transmission Electron Microscopy

The morphology and size of CeO_2_NP were characterized using a Tecnai F20 S/TEM operated in high-resolution mode. For sample preparation, 10 µL of the nanoparticle suspension was drop-cast onto a 200-mesh carbon-coated copper grid and air-dried at room temperature. To ensure complete drying, samples were allowed to dry for a minimum of 24 hours. Nanoparticle size and size distribution were determined using ImageJ software. A minimum of 2000 particles were measured to ensure statistical significance (Supplemental Fig. S1).

##### UV-visible Spectroscopy

UV-visible spectra of the nanoparticles were recorded using a Cary 60 spectrophotometer (Agilent Technologies, USA) within a wavelength range of 250-800 nm. Measurements were performed in 1.5 mL plastic cuvettes.

##### Dynamic Light Scattering and ζ-Potential

Hydrodynamic size and ζ-potential were determined using a Malvern Zetasizer Nano ZS (Malvern Instruments, UK) equipped with a 532 nm laser and a 173° backscatter detector. Measurements were conducted in 1 cm path length cells at 25°C, with three independent replicates per sample.

#### Organ Distribution and Cerium Content Determination

Tissue samples were digested using an Ethos™ Easy microwave digestion system (Milestone). Samples were thawed and then combined with a 1:2 (v/v) nitric acid:water digestion solution. Digestion was performed at 200°C for 90 minutes. Cerium content was quantified by ICP-MS (Agilent 7900 ICP-MS) at the Chemical Analysis Service, UAB, Barcelona.

### Experimental design and *in vivo* exposure to diesel exhaust particles and CeO_2_NP

Seventy-five Sprague-Dawley rats (both sexes, in a 1:1 ratio), aged 6 weeks at the start of the study, were used throughout the experiment. Following a four-week acclimatization period, the rats underwent intratracheal instillations of saline containing or not DEPs (7.5 mg/Kg, 0.375 mL/Kg), for one or three weeks, three times a week. The influence of CeO_2_ nanoparticles on the effects of DEP exposure was analyzed in additional rats weekly injected with CeO_2_NP (0.5 mg/Kg the first week, 0.25 mg/Kg the second and third weeks, and 0.5 mg/Kg one day before sacrifice), beginning at the time of the first DEP exposure (Supplemental Fig. S2A). The pro-arrhythmic effects of DEPs were also assessed in rats with myocardial infarction (i.e., subjected to transient coronary occlusion within the 4 weeks prior to DEP exposure) (Supplemental Fig. S2B). Infarctions were induced as previously described ^34^, and the DEP exposure protocol was performed as described above.

Successful DEP exposure was confirmed in the lungs of all experimental animals by fixation and dehydration in methanol, followed by incubation in a mixture of benzyl alcohol/benzyl benzoate (BA/BB) at a 2:1 ratio for at least one week (Supplemental Fig. S2C). Additionally, part of the lungs was fixed in 4% formaldehyde, embedded in paraffin, and sectioned into 4 μm slices. Samples were stained with picrosirius red (Sigma-Aldrich, MO, USA) and examined under a microscope (Eclipse Ts2R-FL, Nikon, Japan) at 40x magnification (Supplemental Fig. S2C).

### Cardiac Function by Echocardiography

Systolic cardiac function was assessed by transthoracic echocardiography using a Vivid Q portable ultrasound system equipped with a 13 MHz i12L-RS probe (GE Healthcare), as previously described ^34^. Echocardiographic images were acquired at the end of the 3-week exposure period, one day before euthanasia.

### Isolated, Langendorff-perfused, rat heart preparation

Three weeks after DEP or sham exposure, rats (13 weeks of age, males and females weighing 400–500 g and 250–300 g, respectively) were anesthetized with sodium pentobarbital (1.5 g/Kg, IP) and underwent a bilateral thoracotomy. Whole hearts were quickly excised and retrogradely perfused through the aorta with oxygenated (95% O2: 5% CO2) Krebs solution at 37°C (composition in mmol/L: NaCl 118, KCl 4.7, MgSO_4_ 1.2, CaCl_2_ 1.8, NaHCO_3_ 25, KH_2_PO_4_ 1.2, glucose 11, pH 7.4) in a constant flow Langendorff system, as previously described ^35^.

### Electrogram recordings

Electrograms were recorded using stainless steel electrodes (Model 6491 unipolar pediatric temporary pacing lead, Medtronic France, Fourmies, France) placed at the left ventricular base and at the metallic cannula used to perfuse the heart, as previously described ^36^. Changes in the duration of the P wave, PR interval, QRS complex, and Bazett corrected QT interval were analyzed in these recordings.

### Electrophysiological studies

In all experimental groups, spontaneous ventricular arrhythmias were studied during a 10-minute control period. Thereafter, inducible ventricular arrhythmias were triggered using two additional stainless steel electrodes placed at the cardiac apex and a protocol of programmed electrical stimulation, as previously described ^36^. This protocol consisted of a train of 9 stimuli (S1), administered at a basic cycle length (BCL) of 150 ms, followed by 1 to 3 extrastimuli (S2-S4) (Fig. 1A). The number and duration of premature ventricular beats (PVB), non-sustained ventricular tachyarrhythmias (NSVT, defined as those lasting less than 30 seconds), and sustained ventricular tachyarrhythmias (SVT, lasting more than 30 seconds) were recorded during the entire protocol. The number of arrhythmic episodes was expressed as the count of events divided by the total stimulation cycles applied to each heart.

**Figure 1.**
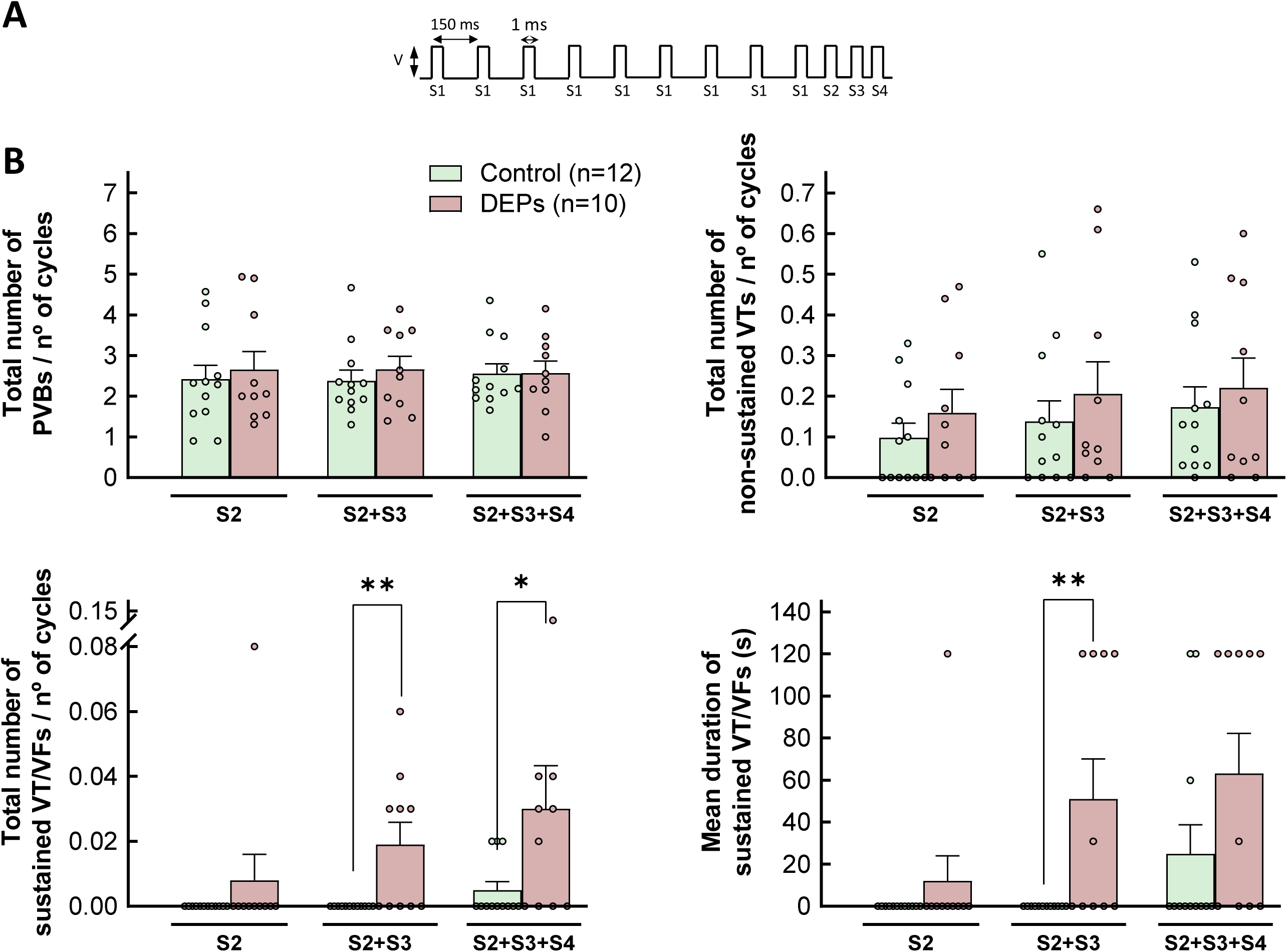
(A) Outline of the protocol of programmed electrical stimulation used in the study to induce ventricular arrhythmias. A train of 9 stimuli (S1), separated 150 ms, was followed by 1 to 3 extrastimuli (S2-S4). Pulse duration was set at 1 ms, and amplitude was set at a voltage (V) double of the diastolic threshold. (B) Number of premature ventricular beats (PVBs), non-sustained tachycardias (VTs) and sustained tachyarrhythmias detected after application of 1, 2 or 3 extrastimuli, expressed relative to the number of stimulation cycles, in isolated rat hearts from animals intratracheally instilled for three weeks with saline containing or not DEPs. Additionally, the mean duration of sustained VTs was also determined (lower, right, panel). * (p<0.05) and ** (p<0.01) indicate significant differences vs. the respective control group.

### Heart weight to tibia length and interstitial collagen deposition

Cardiac hypertrophy was calculated as the ratio of heart weight to tibia length (HW/TL). Interstitial collagen deposition was determined in cardiac slices stained with picrosirius red ^37^. In animals that were submitted to transient coronary occlusion, infarct size was determined by quantifying the area of fibrosis stained with picrosirius red, and expressed as a percentage of the total slice area, normalized by cardiac weight.

### Immunofluorescence analysis of connexin 43 (Cx43) distribution

Cx43 remodeling, including changes in expression and/or distribution, have a major influence on the appearance of cardiac arrhythmias ^38^. Accordingly, we assessed Cx43 distribution by confocal laser scan microscopy in cryosections of cardiac samples from control and DEP-treated animals ^36^. Additionally, Cx43 expression was analyzed by conventional Western blot as described below.

### Inflammatory cell infiltration

Immunohistochemistry was performed in snap-frozen OCT-embedded cardiac sections (4 μm) from additional hearts. Sections were first incubated, overnight (4°C), with a rabbit antibody raised against CD45 (ab10558, dilution 1:500, Abcam). Subsequently, the samples were incubated again for 90 min at room temperature with a biotinylated anti-rabbit secondary antibody (Vector Laboratories, Peterborough, UK). The standard Vectastain avidin-biotin peroxidase complex (ABC; Vector Laboratories, Peterborough, UK) was then applied. Color development was performed using 3,3′-diaminobenzidine (DAB), and the sections were counterstained with hematoxylin before dehydration, clearing, and mounting. Negative controls, in which the primary antibody was omitted, were included to assess non-specific binding. All staining procedures were performed in triplicate.

### Analysis of myocardial oxidative stress

Oxidative stress was evaluated in myocardial samples from these additional animals by measuring the ratio of reduced to oxidized glutathione (GSH/GSSG) and the concentration of malondialdehyde (MDA), as a marker of lipid peroxidation.

Total GSH concentrations and those of the reduced and oxidized (GSSG) fractions were determined spectrophotometrically at 412 nm in myocardial extracts using a modification of the previously described GSH reductase enzymatic method ^39^. Results were expressed as nmoles of GSH per milligram of protein. A decrease in the GSH/GSSG ratio was used as an indicator of enhanced oxidative stress.

Additionally, lipid peroxidation, an indirect indicator of oxidative stress, was assessed by homogenizing myocardial tissue in ice-cold RIPA buffer and measuring MDA levels using a TBARS (TCA Method) Assay Kit (700870, Cayman Chemical), according to manufacturer’s instructions. MDA concentrations were colorimetrically quantified at 430 nm using a multimode reader.

### Real time RT-qPCR

Total cardiac RNA was isolated using the TriPure Isolation Reagent (Roche Diagnostics, Indianapolis, IN) according to the manufacturer’s instructions. DNase I-treated total RNA (1 μg) was reverse-transcribed into cDNA using the High-Capacity cDNA Archive Kit (Applied Biosystems, Foster City, CA) with random hexamers. Quantification of mRNA levels was performed by real-time PCR using an ABI PRISM 7900HT sequence detection system (Applied Biosystems, Foster City, CA) and specific primers and probes for rat provided by Applied Biosystems (Assay-on-Demand system, see the Supplemental material for details). Relative mRNA levels were determined using the 2^−ΔΔCt^ method.

### Western blot analysis

Protein expression was analyzed by conventional Western blot, as previously described ^37^. Extracts were electrophoretically separated on 10-12% polyacrylamide gels and band intensities were measured by densitometry using the Image Studio Lite software.

### Statistics

Data are expressed as mean±SEM. Differences in the number of spontaneous or induced ventricular arrhythmias were assessed using nonparametric Mann-Whitney or Kruskal-Wallis and Dunn’s tests. Additional analyses were performed by two-way ANOVA to determine the effects of previous myocardial infarction and DEP exposure or the impact of sex. Incidences were assessed by χ^2^ or Fisher’s exact tests. Changes in HW/TL, echocardiographic measurements, collagen deposition, inflammatory cell infiltration, RT-PCR, and Western blot data were analyzed using Student’s t-test or one-way ANOVA with Tukey post-hoc tests. Differences were considered significant when p<0.05.

## RESULTS

### Arrhythmia inducibility in isolated rat hearts from animals exposed to DEPs

Spontaneous arrhythmias appeared only as occasional isolated PVB, with no differences between experimental groups (data not shown).

The application of the protocol of programmed electrical stimulation to isolated rat hearts from control animals induced only a few arrhythmic events, consisting of PVB along with some isolated NSVT. Intratracheal exposure to DEPs did not alter the number of either PVB or NSVT (Fig. 1B). In contrast, DEP exposure significantly increased the incidence (0 out of 12 hearts from control rats vs. 5 out of 10 hearts from DEP-exposed animals after application of 2 extrastimuli, Fisher’s exact test, p=0.0096), the number and duration of SVT, effect that was particularly evident with two and three extrastimuli (Fig. 1B). These effects were independent of sex (two-way ANOVA, p-NS). On the other hand, DEP exposure did not modify ventricular refractory periods as compared with control animals (Supplemental Fig. S3A).

In animals with a prior myocardial infarction, later DEP exposure did not influence scar size (Supplemental Fig. S3B). Conversely, a previous myocardial infarction further increased arrhythmogenesis in isolated rat hearts, both in the absence and presence of DEP exposure. Indeed, two-way ANOVA revealed a significant effect of myocardial infarction in the number of both NSVT and SVT following application of 1, 2 or 3 extrastimuli (Supplemental Fig. S3C). Additionally, significant effects on arrhythmia duration were observed for both myocardial infarction and DEP exposure after 1 and 2 extrastimuli (Supplemental Fig. S3C). Further analysis using the non-parametric Krustal-Wallis and Dunn’s test showed a significant effect in the number of SVT and in arrhythmia duration in infarcted animals exposed to DEPs (Supplemental Fig. S3C).

### Effects of DEP exposure on cardiac remodeling and the arrhythmogenic substrate

Given that the major proarrhythmic effects were observed in healthy animals exposed to DEPs, subsequent experiments were performed exclusively in rats with no prior infarction subjected to saline or DEPs instillation. Intratracheal administration of DEPs for three weeks did not alter the HW/TL ratio (Supplemental Fig. S4A). These data were confirmed by echocardiographic analyses, which did not show significant changes on cardiac dimensions and function (Supplemental Fig. S4B). Further, no changes were detected in the expression of the hypertrophic markers MYH7 or NNPA between sham animals and those exposed to DEPs (Supplemental Fig. S4C).

In contrast, three-week exposure to DEPs led to a moderate increase in interstitial collagen deposition (Fig. 2A). This pro-fibrotic response appears to constitute an early response to DEP exposure, as indicated by elevated mRNA levels of the fibrotic markers COL1A1, COL3A1, LOX, LOXL2, and TGFβ1, and of the extracellular matrix-related enzyme MMP2, within one week of intratracheal DEP instillation (Fig. 2B). Remarkably, by three weeks, expression of these markers returned to control levels (Fig. 2B). Western blot analysis revealed a similar pattern for the active form of TGFβ, while α-smooth muscle actin was enhanced at the end of the exposure period (Fig. 2C).

**Figure 2.**
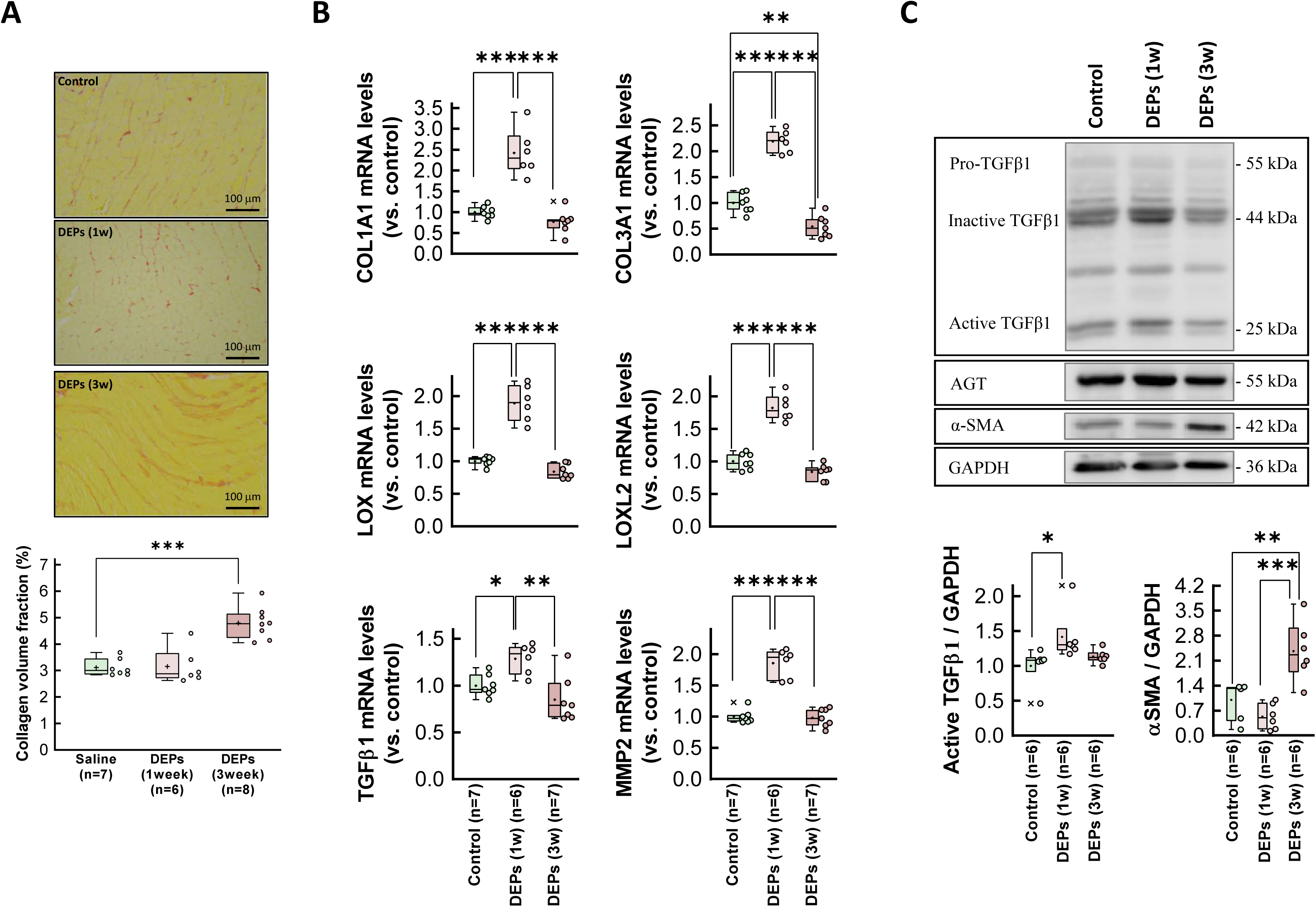
(A) Interstitial collagen deposition in hearts from rats intratracheally instilled with saline containing or not DEPs after 1 or 3 weeks of exposure. (B) Myocardial levels of mRNAs coding for proteins involved in the fibrotic response (COL1A1, COL3A1, LOX, LOXL2, TGFβ1 and MMP2) analyzed in tissue extracts from the same experimental groups. Values are expressed as fold change with respect to rat adenine phosphoribosyltransferase (APRT), used as a housekeeping gene, and normalized respect to control hearts. (C) Representative Western blot analysis showing expression of TGFβ1, NF-κB, angiotensin receptor (AGT), αSMA and GAPDH, in the same experimental groups. Quantifications of active TGFβ and αSMA are shown below. Data are shown as box plot depicting median (horizontal line), mean (+), and individual values (color symbols). Outliers are marked with “x”, and were also included in the analyses. * (p<0.05), ** (p<0.01) and *** (p<0.001) indicate significant differences vs. indicated groups.

Western blot analysis of Cx43 demonstrated three specific immunoreactive bands appearing between 41 and 43 kDa, corresponding to different phosphorylation states ^38^, in myocardial samples from rat hearts. Quantification of these bands did not reveal any difference between control and DEP-exposed animals (Supplemental Fig. S5A). Additionally, confocal analysis did not demonstrate any alteration in Cx43 distribution, with most distributed within the intercalated discs, and presenting only minor lateralization, which was similar in all experimental groups (Supplemental Fig. S5B).

Figure 3A shows a representative cardiac electrogram from an isolated heart included in the control group of animals. Intratracheal instillation of DEPs for three weeks resulted in a prolongation of the P wave and the QRS complex, along with an increased duration of the corrected QT interval (Fig. 3). In contrast, no changes were observed in the duration of the PR interval.

**Figure 3.**
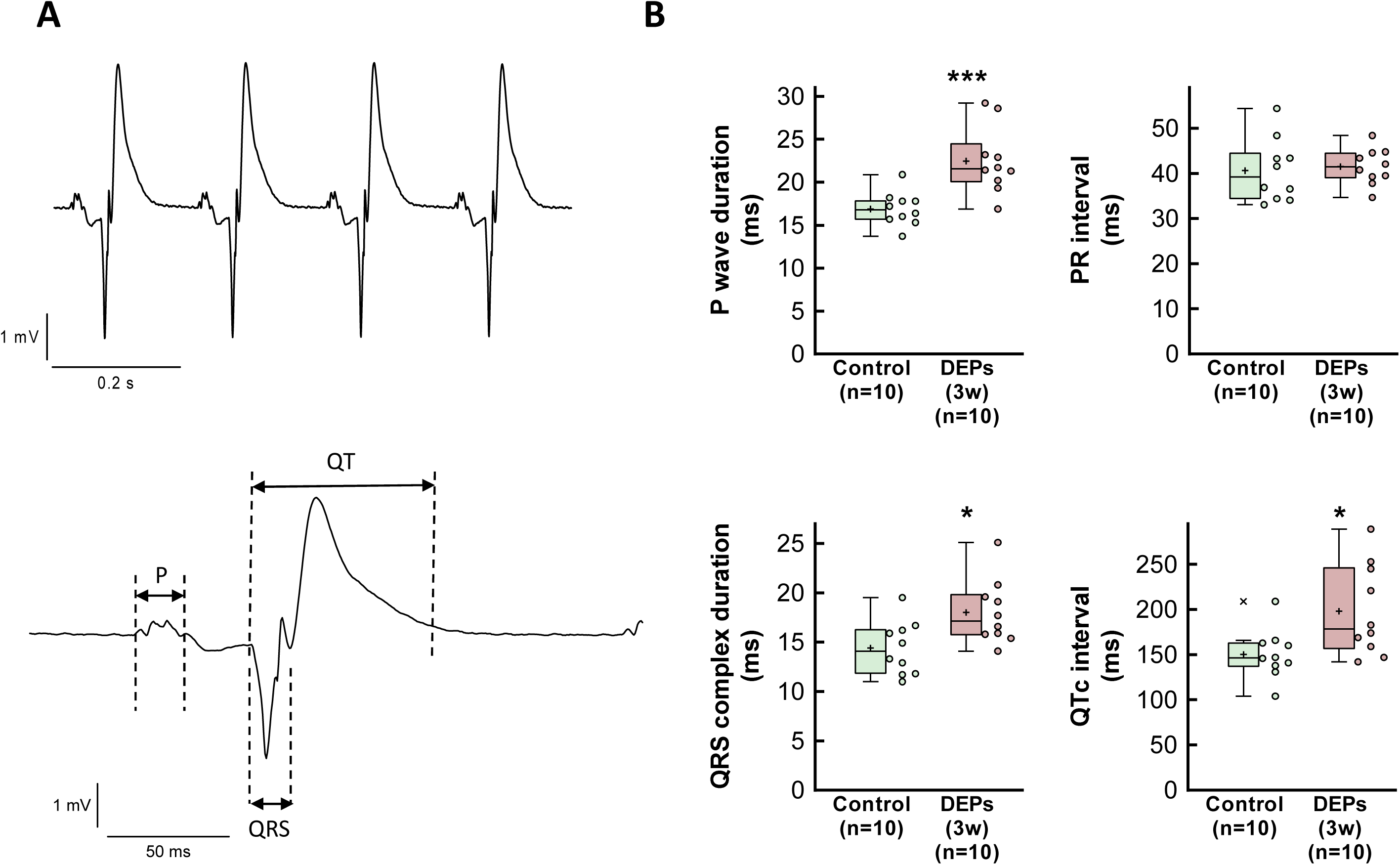
(A) Representative electrogram recording from a heart from a control animal included in the study. The main waves and complexes are shown below. (B) Electrogram characteristics in isolated rat hearts from animals intratracheally instilled for three weeks with saline containing or not DEPs. Data are shown as box plot depicting median (horizontal line), mean (+), and individual values (color symbols). * (p<0.05) and *** (p<0.001) indicate significant differences vs. control hearts.

### Effects of DEP exposure on cardiac inflammation

Immunohistochemistry using an anti-CD45 antibody revealed an increased inflammatory cell infiltrate in myocardial samples from DEP-exposed rats (Fig. 4A). Further, Western blot analysis demonstrated a significant early increase in NF-κB p65, a transcription factor involved in the development and progression of inflammation in the cardiovascular system ^40^, following intratracheal DEP instillation, with partial recovery at three weeks (Fig. 4B). Consistently with the presence of myocardial inflammation, RT-qPCR analysis showed elevated mRNA levels of IL-1β and IL6 shortly after DEP exposure, with a similar trend for TNFα and the macrophage marker EMR1 (Fig. 4C). Notably, myocardial levels of these inflammatory markers returned to baseline three weeks after DEP exposure.

**Figure 4.**
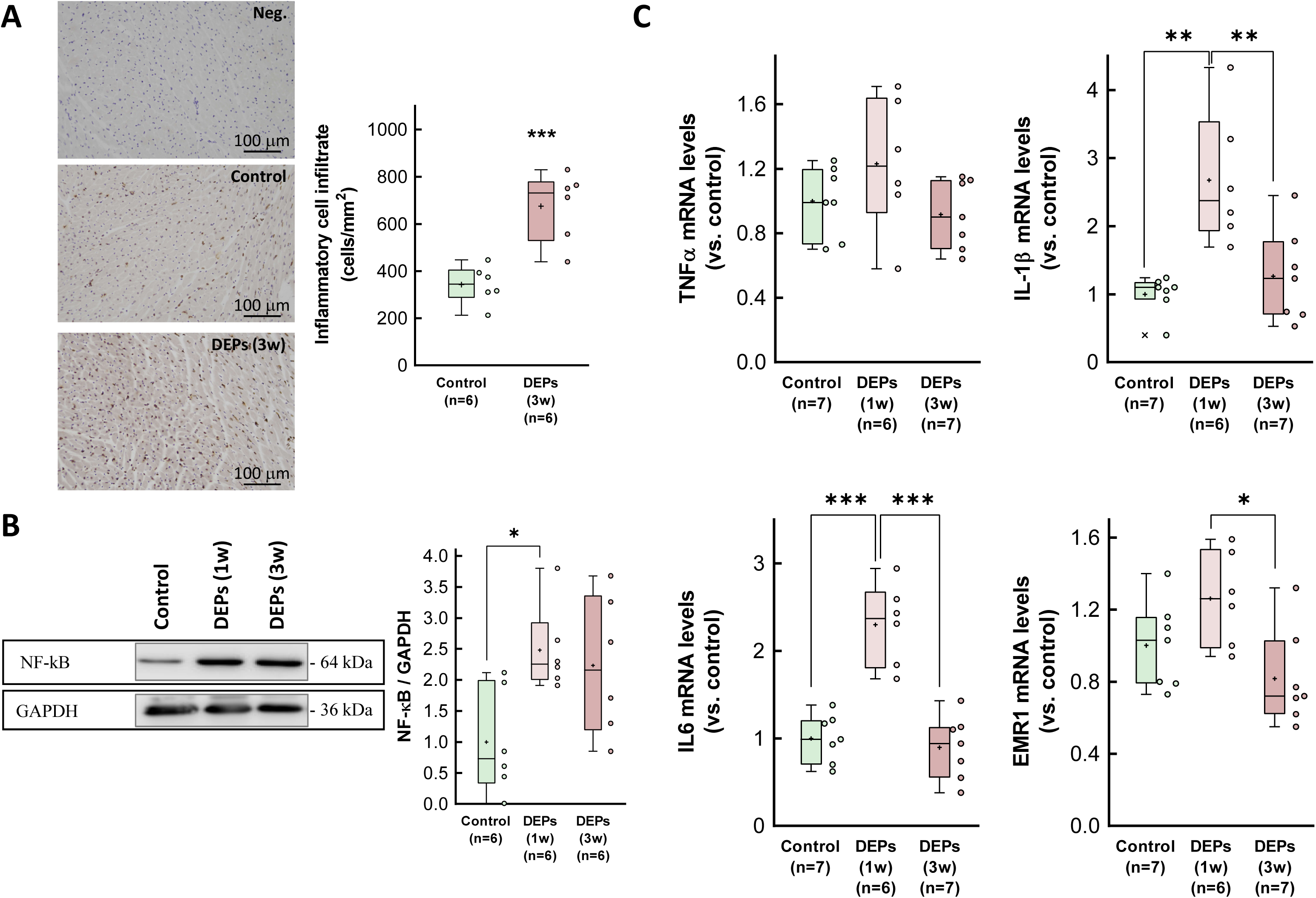
(A) Representative immunohistochemical images of cardiac slices obtained from a control rat and a DEP-instilled animal, incubated with antibodies raised against CD45, and showing intense immune cell infiltration after three weeks of exposure. (B) Expression of NF-κB p65, assessed by Western blot, in myocardial samples from control animals or from rats exposed to DEPs for 1 or 3 weeks. (C) Myocardial levels of mRNAs coding for proteins involved in the inflammatory response (TNFα, IL-1β, IL6 and EMR1) analyzed in tissue extracts from the same experimental groups. Values are expressed as fold change with respect to rat adenine phosphoribosyltransferase (APRT), used as a housekeeping gene, and normalized respect to control hearts. Data are shown as box plot depicting median (horizontal line), mean (+), and individual values (color symbols). Outliers are marked with “x”, and were also included in the analyses. * (p<0.05), ** (p<0.01) and *** (p<0.001) show significant differences vs. indicated groups.

### Myocardial oxidative stress after DEP exposure

The ratio of reduced to oxidized gluthatione (GSH/GSSG) was significantly diminished in myocardial samples from animals intratracheally exposed to DEPs for three weeks, suggesting enhanced oxidative stress (Fig. 5A). Consistently, measurement of myocardial MDA concentrations, a marker of lipid peroxidation, were enhanced at the end of the exposure period (Fig. 5B).

**Figure 5.**
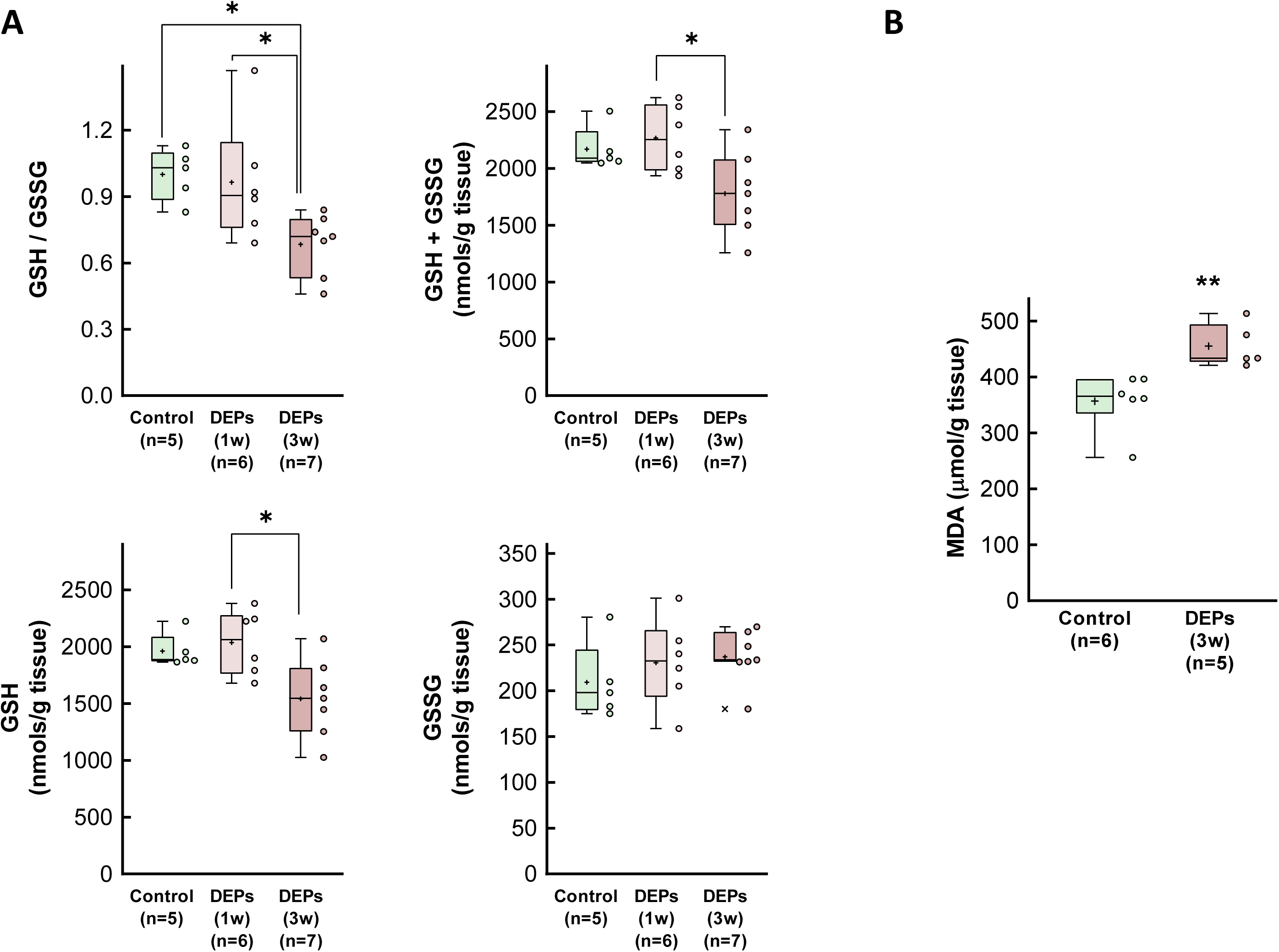
(A) Reduced to oxidized glutathione (GSH/GSSG) ratio, together with total GSH concentrations as well as those of the oxidized (GSSG) and reduced (GSH) fractions in myocardial samples from control animals and from rats exposed to DEPs for 1 or 3 weeks. (B) MDA concentrations in myocardial samples from control rats and from animals exposed to DEPs for 3 weeks. Data are shown as box plot depicting median (horizontal line), mean (+), and individual values (color symbols). Outliers are marked with “x”, and were also included in the analyses. * (p<0.05) and ** (p<0.01) show significant differences vs. indicated groups.

### Activation of cytosolic signaling pathways

Previous studies have linked the effects of PM to the activation of various cytosolic signaling proteins, including ERK1/2, p38 MAPK and Akt ^41,42^. To determine whether the pro-arrhythmic effects of DEP exposure were associated with the activation of these kinases, we assessed their activation and expression by Western blot analysis. Our results showed increased activation (i.e., phosphorylation) of ERK1/2 and decreased phosphorylation of GSK3β one week after intratracheal instillation. No changes were observed in other cytosolic signaling molecules, including Akt, p38 MAPK, SMAD2/3 or TAK1 (Fig. 6 and Supplemental Fig. S6). In all cases, total protein levels remained unchanged.

**Figure 6.**
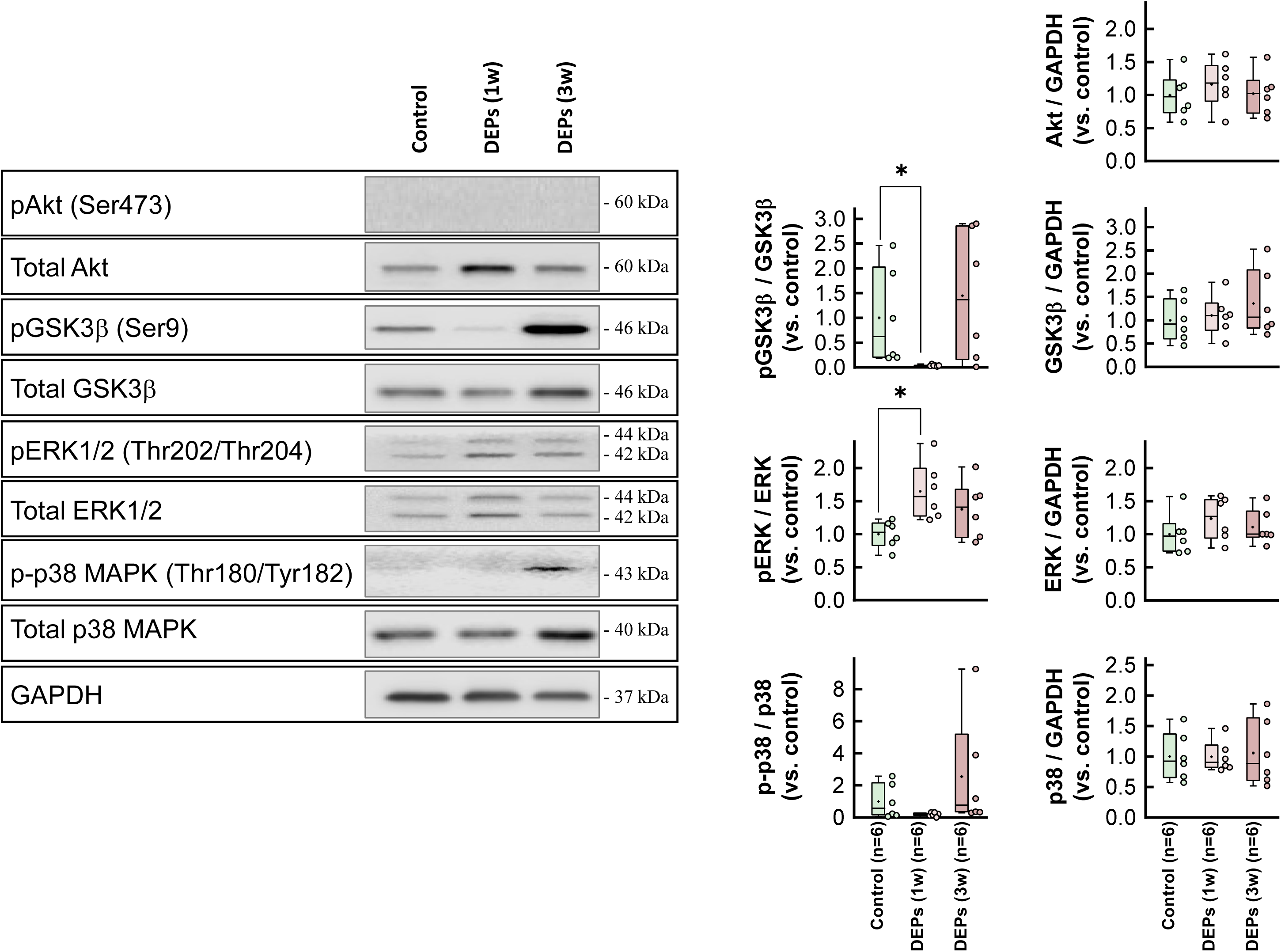
Representative Western blot analysis showing expression and degree of activation (i.e., phosphorylation) of Akt, GSK3β, ERK1/2 and p38 MAPK in myocardial samples from control animals and from rats exposed to DEPs for 1 or 3 weeks. Quantifications are shown at the right. No activation was detected for Akt in any case. Data are shown as box plot depicting median (horizontal line), mean (+), and individual values (color symbols). * (p<0.05) show significant differences vs. indicated groups.

### Effects of CeO_2_ nanoparticles on the pro-arrhythmic impact of DEPs

Our findings confirm that DEP exposure is associated with increased arrhythmia vulnerability in healthy hearts, with several potential underlying mechanisms, of which only an increase in oxidative stress remains activated in the long term. Therefore, we next assessed whether treatment with CeO_2_NP, a ROS-detoxifying strategy, could mitigate the effects of DEP exposure on arrhythmia susceptibility.

Weekly treatment with a CeO_2_NP solution of 1 mg/mL (stabilized with 10 mg/mL RSA in 10 mmol/L phosphate buffer, at 0.5 mg/Kg the first week, 0.25 mg/Kg the second and third weeks, and 0.5 mg/Kg one day before sacrifice) in rats subjected to DEP exposure resulted in a trend towards a reduction in NSVT and led to a significant, complete abolition of SVT compared to DEP-treated rats and no CeO_2_NPs treatment (0 out of 9 animals, p=0.0325 vs. DEP-treated rats, Fisher’s exact test) (Fig. 7). The anti-arrhythmic effects of CeO_2_NPs were associated with reversion of the ECG changes induced by DEP (Fig. 8A) and with restoration of interstitial collagen deposition (Fig. 8B), MDA concentration (Fig. 8C), and immune cell infiltration (Fig. 8D) to control levels. Further, RT-qPCR analysis confirmed a reduction in the expression of most fibrotic markers one week after DEP exposure (Fig. 9A) along with decreased IL6, though IL-1β mRNA levels remained unchanged (Fig. 9B). Surprisingly, however, CeO_2_NPs treatment was associated with enhanced levels of TNFα mRNA levels (Fig. 9B). Remarkably, cerium, detected by ICP-MS, was only present in the liver and the spleen of these animals, with only residual amounts in the heart or other tissues (Supplemental Fig. S7).

**Figure 7.**
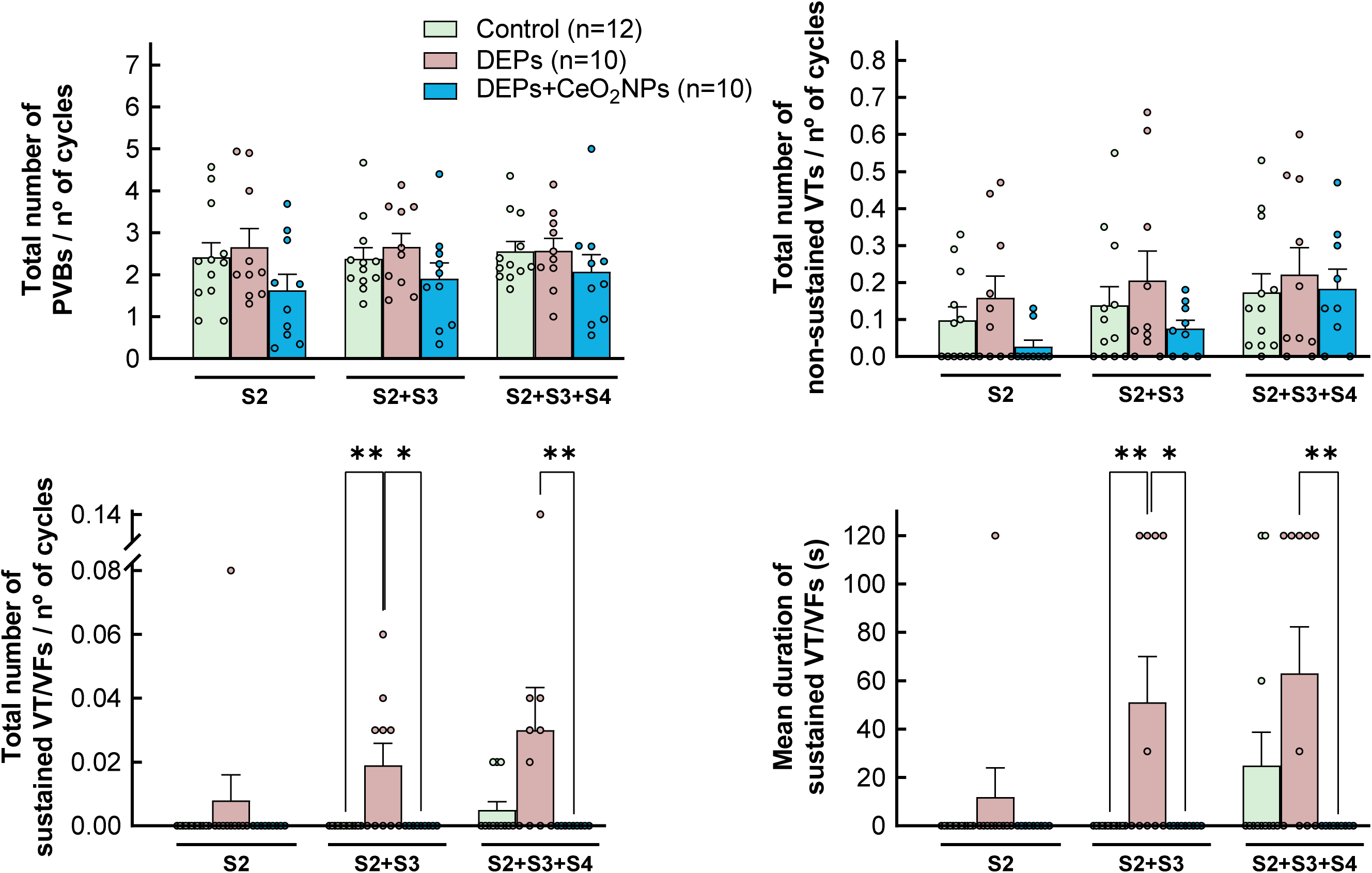
Number of premature ventricular beats (PVBs), non-sustained tachycardias (VTs) and sustained tachyarrhythmias detected after application of 1, 2 or 3 extrastimuli, expressed relative to the number of stimulation cycles, in isolated rat hearts from animals intratracheally instilled for three weeks with saline or DEPs, either with or without additional injection of CeO_2_NP. Mean duration of sustained tachyarrhythmias is shown at the right bottom. * (p<0.05) and ** (p<0.01) show significant differences vs. indicated groups.

**Figure 8.**
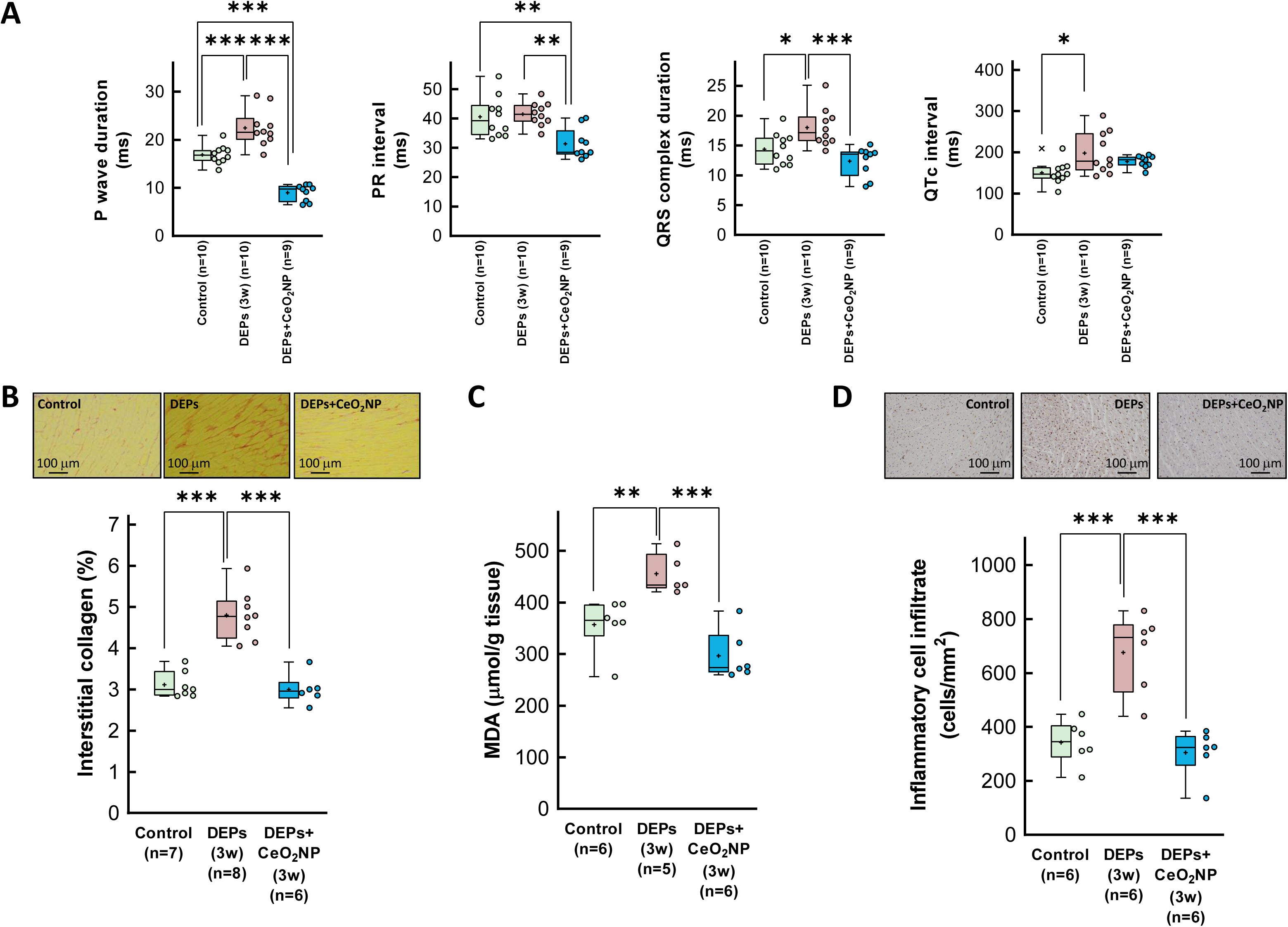
Electrogram characteristics (A), interstitial collagen deposition (B), MDA concentrations (C) and immune cell infiltration (D) in isolated rat hearts from animals intratracheally instilled for three weeks with saline containing or not DEPs, injected or not with CeO_2_NP. Data are shown as box plot depicting median (horizontal line), mean (+), and individual values (color symbols). Outliers are marked with “x”, and were also included in the analyses. * (p<0.05), ** (p<0.01) and *** (p<0.001) show significant differences vs. indicated groups.

**Figure 9.**
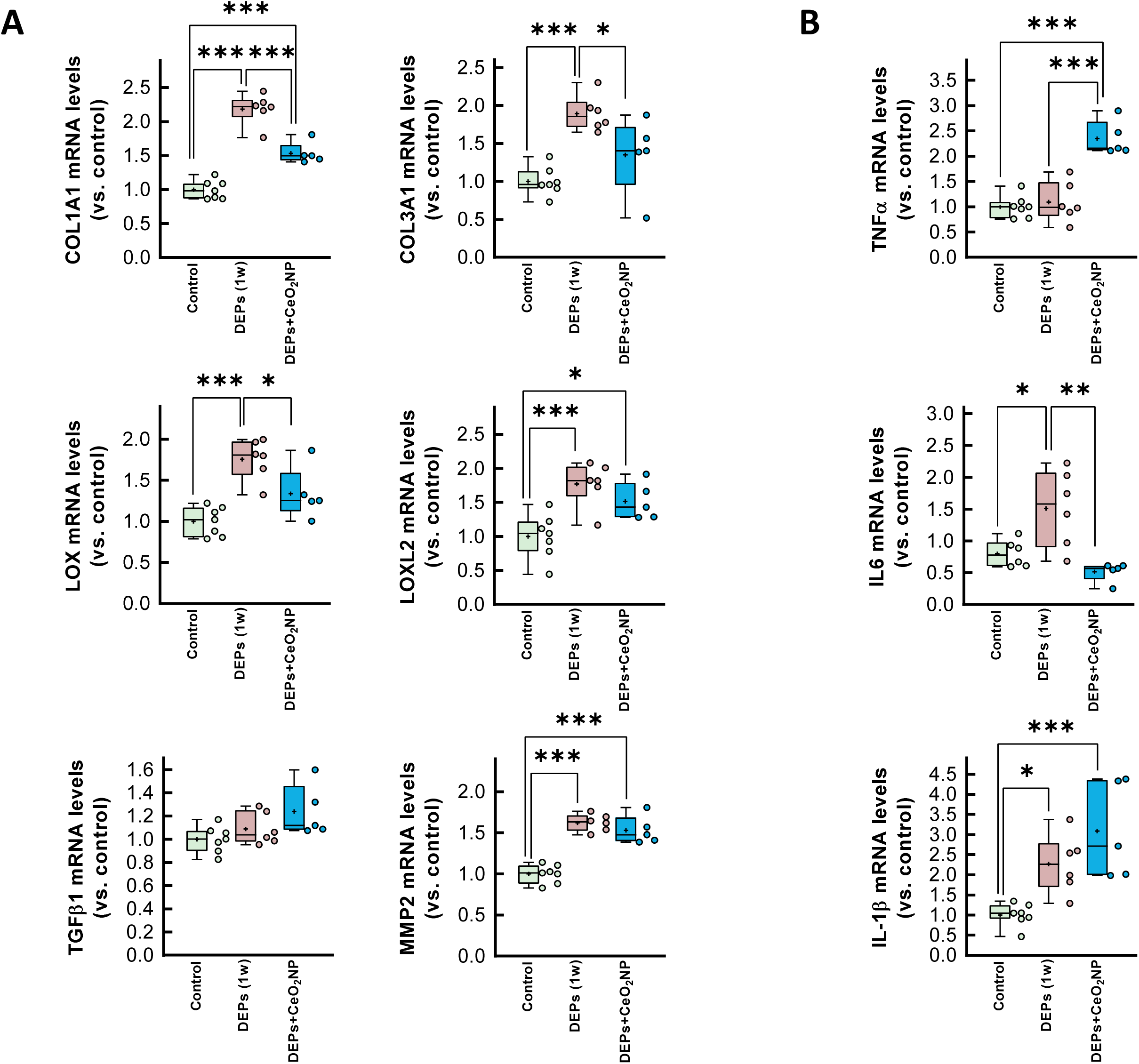
Myocardial levels of mRNAs coding for proteins involved in the fibrotic (A) (COL1A1, COL3A1, LOX, LOXL2, TGFβ1 and MMP2) and inflammatory (B) (TNFα, IL-1β and IL6) responses, analyzed in cardiac extracts from animals intratracheally instilled for three weeks with saline containing or not DEPs, injected or not with CeO_2_NP. Data are shown as box plot depicting median (horizontal line), mean (+), and individual values (color symbols). * (p<0.05), ** (p<0.01) and *** (p<0.001) show significant differences vs. indicated groups.

## DISCUSSION

This study demonstrates that a three-week exposure to DEPs induces a pro-arrhythmic effect in isolated rat hearts. This effect was associated with a moderate increase in interstitial collagen deposition, ECG alterations, enhanced myocardial inflammation, sustained increased oxidative stress, and activation of specific cytosolic signaling pathways. Notably, treatment with CeO_2_NPs, a known ROS scavenger, attenuates the pro-arrhythmic effects of DEPs, reduces myocardial fibrosis and oxidative stress, and reverses most of these changes.

Evidences linking air pollution to the occurrence of cardiac arrhythmias remains controversial ^8^. Clinical and epidemiological studies have yielded contradictory results. In a cohort of patients with implanted cardioverter defibrillators from the Boston metropolitan area, followed for 3.1 years, no statistically significant associations were observed between ventricular arrhythmic episodes and any analyzed pollutant, including PM2.5 on the same or previous days ^16^. Similarly, a study in Sweden found no positive association between atrial fibrillation and long-term residential air pollution exposures ^17^. In contrast, other studies have demonstrated an increased risk of cardiac arrhythmias following both short- ^18–20^ and long-term ^21–23^ exposure to PM.

Most experimental studies have used DEPs as a surrogate for air pollution to assess its impact on cardiac arrhythmias. Previous short-term exposure to DEPs was shown to enhance ventricular arrhythmia duration during ischemia in rats submitted to transient coronary occlusion followed by reperfusion ^24^. Furthermore, a single exposure to particulate matter was shown to elevate the risk of aconitine-induced cardiac arrhythmias, including premature ventricular beats and ventricular tachyarrhythmias, in hypertensive rats ^26,27^. However, these studies were conducted under conditions that predispose to arrhythmias. To our knowledge, a single group has investigated the effects of short-term DEP exposure on cardiac arrhythmias in healthy rats, reporting an increased incidence of spontaneous triggered activities and VT 30 min after intratracheal instillation ^29,30^. Our present data demonstrate that DEP exposure not only enhances short-term ventricular arrhythmogenesis but also induces pro-arrhythmic effects after three weeks of chronic exposure in isolated hearts from healthy rats. Specifically, we observed an increase in both the incidence and duration of SVT, the most severe arrhythmic events, which may potentially compromise patient survival. Additionally, prior myocardial infarction further enhanced arrhythmogenesis in isolated rat hearts, effects that tended to be exacerbated by DEP exposure.

While no changes in cardiac dimensions or Cx43 expression and distribution were observed between control and DEP-exposed hearts, the pro-arrhythmic effects detected three weeks post-instillation were associated, in our study, with a moderate but significant increase in myocardial fibrosis. Interstitial collagen deposition appears to be an early response, as evidenced by the elevated mRNA levels of fibrosis-associated markers, including COL1A1, COL3A1, and LOX, and of the extracellular matrix-related enzyme MMP2, just one week after exposure. Similarly, both mRNA and protein levels of the active form of TGFβ were upregulated soon after DEP exposure. Notably, these transcriptional changes returned to baseline by three weeks, suggesting an adaptative response. This pro-fibrotic response observed by us and others ^42–44^ may contribute to the pro-arrhythmic effects of DEPs. Indeed, previous studies have demonstrated that myocardial fibrosis predisposes individuals to cardiac arrhythmias by altering anisotropic conduction, inducing source-to-sink mismatch, and enhancing refractoriness dispersion^45–47^.

The enhanced fibrotic response may underlie some of the ECG changes we have observed after DEP exposure. Increased P wave duration and dispersion have been correlated with enhanced atrial fibrosis ^48^, while QRS duration is similarly associated with ventricular fibrosis ^49^. Fibrosis can potentially reduce conduction velocity in both cardiac chambers, leading to the prolongation of these two ECG parameters. Additionally, we observed an increased QTc complex duration (that is primarily determined by the action potential duration ^50^), which alone may contribute to the development of cardiac arrhythmias. QT complex prolongation can lead to early afterdepolarizations, which, in turn, may trigger ventricular arrhythmias, particularly torsades de pointes (TdP), a potentially life-threatening form of polymorphic ventricular tachycardia ^51^. Our findings align with previous studies demonstrating prolongation of the QRS complex ^27,43^ and of either the QT interval ^27,29,43^ or action potential duration ^29,30^ after exposure to DEPs, synthetic residual oil fly ash particles, or particulate matter from ambient air.

Upon inhalation, PM is rapidly phagocytosed by alveolar macrophages, which become over-activated triggering an inflammatory response in the lung ^52^. Neutrophil and macrophage activation potentiate cytokine secretion ^53^ and further recruits additional inflammatory cells, thereby amplifying tissue injury and contributing to systemic and cardiovascular inflammation ^52,54^. Our present findings demonstrate enhanced cardiac inflammation, as demonstrated by an increased inflammatory cell infiltrate in the myocardium. Myocardial inflammation appears to constitute an early response to DEP exposure, paralleling the expression of fibrotic markers. This is supported by elevated

NF-κB in Western blot analysis and increased mRNA levels of IL6 and IL-1β one week after DEP exposure, with similar trends for TNFα and EMR1. However, similar to fibrosis, an adaptative response seems to occur by three weeks, as these mRNA levels return to baseline.

Inflammation goes hand-in-hand with oxidative stress, with inflammatory cells producing ROS, while ROS, in turn, enhance inflammation through NLRP3 inflammasome activation ^55^, creating a positive feedback that exacerbates tissue injury ^54^. Our current findings demonstrate a decreased GSH/GSSG ratio three weeks after exposure, accompanied by increased myocardial MDA concentrations, both of which are indicative of heightened oxidative stress and lipid peroxidation. Notably, increased oxidative stress may have significant consequences in sarcolemmal currents that may favor the appearance of arrhythmias ^56^. In this sense, increased oxidative stress has been shown to enhance the slowly inactivating component of the sodium current (late I_Na_), leading to prolongation of both cardiomyocyte action potentials and QT interval and appearance of early afterdepolarizations ^57^. Further, ROS have been shown to promote fibroblast differentiation into myofibroblasts ^58^, contributing to interstitial fibrosis, as suggested in our study by elevated αSMA levels. In this regard, we observed enhanced ERK1/2 activation in myocardial samples from DEP-exposed animals, supporting previous studies implicating the AngII/ERK1/2/TGFβ1 signaling pathway in PM2.5-induced fibrosis ^42^. Furthermore, our Western blot analysis revealed decreased phosphorylation (i.e., enhanced activation) of GSK3β one week after intratracheal instillation. Given GSK3β‘s critical role in mitochondrial regulation ^59^, this finding aligns with previous reports of mitochondrial dysfunction following DEP exposure ^60^.

Notably, our findings reveal that the pro-arrhythmic effects of DEPs were reversed by treatment with CeO_2_NPs. Indeed, CeO_2_NPs have been shown to confer cardioprotective effects against several ROS-dependent pathologies. They attenuated QT prolongation, cardiac enzyme release, oxidative stress and apoptosis in a rat model of doxorubicin-induced cardiomyopathy ^61^. Similarly, CeO_2_NPs reduced left ventricular dysfunction and dilatation in mice with cardiac-specific expression of monocyte chemoattractant protein (MCP)-1 expression ^62^, and mitigated cardiomyopathy secondary to obesity induced by a high-fat, high sucrose, diet ^63^. Additionally, CeO_2_NPs diminished monocrotaline-induced pulmonary hypertension and associated right ventricular hypertrophy in rats ^64^. Finally, CeO_2_NPs-decorated polycaprolactone (PCL)-gelatin blend nanofibers significantly reduced ROS levels and attenuated agonist-induced hypertrophy in neonatal rat primary cardiomyocytes ^65^. To our knowledge, this is the first study to analyze the effects of CeO_2_NPs on ventricular arrhythmogenesis.

The antiarrhythmic effects of CeO_2_NPs were associated with a reduction in oxidative stress, as indicated by the recovery in MDA levels. This finding underscores not only the potential of CeO_2_NPs as a therapeutic strategy but also the critical role of ROS in the harmful cardiovascular effects of air pollution, particularly DEPs. This aligns with previous studies suggesting that antioxidants such as N-acetylcysteine (NAC) may mitigate the adverse cardiac effects of PM ^29,66^. However, the use of NAC as a safe antioxidant has come under debate due to a lack of evidence regarding its antioxidant effects and its poor pharmacology ^67^. As expected, the reduction in oxidative stress was accompanied by decreased collagen deposition three weeks after exposure, downregulation of fibrosis-related mRNA markers, a decrease in infiltrating inflammatory cells, shortened P wave, QRS and QTc durations, and attenuated IL6 expression. Contrarily, however, CeO_2_NPs-treated hearts showed an increase in TNFα mRNA levels. The underlying mechanism for this discrepancy remains unclear, but it may suggest that CeO_2_NPs also induce some degree of immune response.

As previously described, iv administered CeO_2_NPs majorly is found in the liver and spleen ^68,69^, as we observe here, where they are well tolerated ^33^. A small fraction is found in the lungs one day after the last administration. CeO_2_NPs are known to progressively accumulate in the liver and spleen and, from there, be slowly excreted through the hepatobiliary route ^68,70^; therefore, it is possible that lung concentrations are more significant at earlier times. It is worth noting that the ROS scavenging and therefore anti-inflammatory activity of ceria is intrinsically catalytic, this is, CeO_2_NPs bring together free radicals to force their recombination and are not consumed during the reaction, therefore even at low doses, their ROS scavenging action is permanent allowing for large effects at normally low doses ^71^. Besides, DEPs are known to trigger inflammation in the lungs, potentially leading to systemic inflammation that can damage the cardiovascular system. Likewise, in the case of inflammation becoming systemic, CeO_2_NPs can carry on their action remotely from the liver and the spleen, possibly by limiting systemic inflammation and ROS production and propagation. However, how this is translated into reduced myocardial inflammation and oxidative stress remains to be further investigated.

In conclusion, prolonged exposure to DEPs has significant cardiac consequences, including pro-arrhythmic effects, myocardial inflammation, oxidative stress, and fibrosis, effects that were attenuated by CeO_2_NPs, highlighting their potential as a therapeutic intervention. These findings highlight the crucial role of oxidative stress in DEP-related cardiac damage, and emphasize the impact of nanotherapeutics for preventing the cardiovascular complications associated with air pollution exposure.

## LIST OF ABBREVIATIONS

BA/BB: benzyl alcohol/benzyl benzoate
BCL: basic cycle length
CeO_2_NP: cerium oxide nanoparticles
Cx43: connexin 43
DEP: diesel exhaust particles
GSH: reduced glutathione
GSSG: oxidized glutathione
HW/TL: heart weight to tibia length
MDA: malondialdehyde
NAC: N-acetylcysteine
NSVT: non-sustained ventricular tachycardia
PM: particulate matter
PVB: premature ventricular beats
ROS: reactive oxygen species
RSA: rat serum albumin
SVT: sustained ventricular tachyarrhythmia

## ACKNOWLEDGEMENTS.

We thank Silvia Aguiló for her technical assistance.

## SUPPLEMENTAL MATERIAL

Supplemental methods

Figures S1-S6

References #34 - #38

